# LightMed: A Light-weight and Robust FFT-Based Model for Adversarially Resilient Medical Image Segmentation

**DOI:** 10.1101/2024.09.28.615584

**Authors:** Viet Tien Pham, Minh Hieu Ha, Bao V. Q. Bui, Truong Son Hy

## Abstract

Accurate and reliable medical image segmentation is essential for computer-aided diagnosis and formulating appropriate treatment plans. However, real-world challenges such as suboptimal image quality and computational resource constraints hinder the effective deployment of deep learning-based segmentation models. To address these issues, we propose LightMed, a novel efficient neural architecture based on Fast Fourier Transform (FFT). Different from prior works, our model directly learns on the frequency domain, harnessing its resilience to noise and uneven brightness, which common artifacts found in medical images. By focusing on low-frequency image components, we significantly reduce computational complexity while preserving essential image features. Our deep learning architecture extracts discriminative features directly from the Fourier domain, leading to improved segmentation accuracy and robustness compared to traditional spatial domain methods. Additionally, we propose a new benchmark incorporating various levels of Gaussian noise to assess susceptibility to noise attacks. The experimental results demonstrate that LightMed not only effectively eliminates noise and consistently achieves accurate image segmentation but also shows robust resistance to imperceptible adversarial attacks compared to other baseline models. Our new benchmark datasets and source code are publicly available at https://github.com/HySonLab/LightMed.

## I. Introduction

**M**EDICAL image segmentation is a fundamental process in medical imaging that involves partitioning an image into distinct regions to enhance the visibility of anatomical or pathological structures. This technique is crucial for accurately identifying and analyzing various structures within medical images, such as organs, tissues, and lesions. Various imaging modalities, including X-ray, Positron Emission Tomography (PET), Computed Tomography (CT), Magnetic Resonance Imaging (MRI), and Ultrasound (US), are widely used to capture data. Each modality provides unique insights into the body’s internal structures and functions, enabling thorough and precise diagnostic evaluations.

Early traditional approaches to medical image segmentation primarily focused on techniques such as edge detection, template matching techniques, statistical shape models, machine learning [26]–[29]. The use of deep learning for image segmentation has become increasingly prevalent, as it effectively achieves hierarchical feature representation of images. U-Net [3] is one of the pioneering network architectures in the medical image domain, which relies on convolutional neural networks (CNNs) and fully convolutional networks (FCNs). Numerous improved variants inspried by this architecture [4], [5], [9], [12] have been introduced to further enhance segmentation efficiency. Despite its excellent performance, architectures solely based on CNNs often struggle to capture global and long-range semantic interactions. To address this issue, Transformers became a preferential choice for modelling long-range dependencies in computer vision, inspired by success in the nature language processing domain [13]. Vision Transformer is the first work to demonstrate how Transformers can entirely replace standard convolutions in deep neural networks, achieved comparable performance with the CNN-based methods. Liu et al. [16] proposed Swin Transformer, a hierarchical Transformer whose representation is computed with Shifted Windows. Attention-Unet [14], TransUNet [15], and SwinUNet [17] represent notable advancements in cooperating Transformer-based architectures with U-Net framework.

Recent approaches in medical image segmentation have primarily focused on improving accuracy by increasing the complexity of neural network architectures. However, these models have high computational requirements, which can be problematic in resource-constrained medical environments. Transformer-based models [14]–[17], despite achieving state-of-the-art results on several benchmarks, are known for their resource-intensive self-attention mechanisms, which often require specialized hardware for efficient training and inference. Other enhanced variants based on U-shaped architecture [4], [5], [9], [12], which incorporate additional layers such as squeeze-and-excite blocks and atrous spatial pyramid pooling, also significantly increase the number of parameters and computational complexity. Additionally, noise susceptibility is a significant challenge, often caused by sensor malfunctions and other hardware issues in hospital settings. Once processing new noise images, these prior approaches also experience a significant drop in performance. Moreover, these models still suffer from adversarial examples [39], [51], [52], [54], which can cause catastrophic disruptions. These deceptive tricks can be obtained by adding a visually imperceptible perturbation to human beings.

To remedy the issues mentioned, we propose LightMed, a novel and efficient architecture that operates directly in the frequency domain, conducted based on Fast Fourier Transform (FFT). The Fourier transform is a prevalent techniques in applied mathematics, particularly in fields such as signal processing and data compression, based on trigonometric functions [36]. By leveraging the conjugate symmetry properties of Discrete Fourier Transforms [23], we significantly reduce computational complexity while preserving essential image features through focusing only on low-frequency image components. The Fourier representation is also effective at suppressing noise and correcting uneven brightness, which are common artifacts in medical images [36], [37]. Specifically, our model, designed based on U-shaped architecture, first performs a 2D Fourier transformation of the spatial domain to convert them to the frequency domain, followed by discarding the high frequencies in the first half due to the connjugate symmetry, inspried by [23]. LightMed includes a down-sampling encoder and an up-sampling decoder, compassed of specialized Complex Conv-Block and Complex Convolutional Block Attention Module (CCBAM), designed to handle the real and imaginary components of the image’s frequency domain. Taking the aforementioned Fourier domain as input, the network encoder learns meaningful feature representation, followed by reconstruction of decoder. Finally, an inverse 2D Fourier transform is applied to revert the data back to the spatial domain.

To demonstrate the enhanced noise resistance of our model, we introduce a new benchmark dataset with varying levels of Gaussian noise. This dataset is used exclusively during the inference phase to fairly evaluate the noise suppression capabilities of LightMed in comparison to other baseline models. Our experiments show that LightMed not only achieves competitive segmentation performance while significantly reducing computational overhead but also exhibits robust resistance to adversarial attacks compared to state-of-the-art models.

In summary, our contributions can be summarized as:

- We propose LightMed, an efficient and lightweight neural U-shaped architecture operates directly on the Fourier domain. This architecture leverages Complex Conv-Blocks and Complex Convolutional Block Attention Modules (CCBAM), which effectively harness the unique properties of the frequency domain to enhance feature extraction and representation.
- We introduce a new benchmark incorporating various levels of Gaussian noise to assess susceptibility to noise attacks. Our methods in achieving state-of-the-art performance with significantly lower resource requirements.
- We conduct an extensive experimental study to evaluate the capability of LightMed to defend against crafted adversarial examples. The experiments demonstrate that our model is robust against a wide range of adversarial attacks in comparision with other baselines.

## II. Related work

### A. Segmentation on Medical Image

Medical image segmentation plays a crucial role in the advancement of healthcare systems. This process involves dividing medical images into meaningful regions, which is particularly important for accurate disease diagnosis and effective treatment planning. By isolating specific areas of interest within an image, medical professionals can better identify abnormalities, monitor disease progression, and develop tailored treatment planning. A popular deep learning architecture in the field of semantic segmentation for biomedical application is U-Net [3], consists of a contracting path which capture contextual information and reduce the spatial resolution of the input, and an expansive path decode the encoded data via skip connections.

Inspired by UNet architecture [3], Zhou et al. [4] proposed Unet++, where the encoder and decoder subnetworks are connected through a series of nested, dense skip pathways. [5] introduced ResUNet-a architecture, uses a UNet encoder/decoder backbone, in combination with residual connections [6], atrous convolutions [7], pyramid scene parsing pooling [8]. They also offer a variant of the Dice loss function that enhances the convergence speed of semantic segmentation tasks and boosts overall performance, behaves well even when there is a large class imbalance. Jha et al. [9] proposed ResUNet++, which is an enhanced version of standard ResUNet by integrating an additional layer such as squeeze-and-excite block, ASPP, and attention block to the network. DoubleU-Net [12] employs two U-Net architectures in sequence, featuring two encoders and two decoders. The first encoder in this network is a pretrained VGG-19 [10]-a lightweight model as compared to other pretrained models, which has been trained on the ImageNet [11].

Recently, Transformers became a preferred choice for modelling long-range dependencies in computer vision, inspired by their success with self-attention mechanism in natural language processing [13]. Oktay et al. [14] proposed novel self-attention gating module that can be utilised in CNN based standard image analysis models for dense label predictions. In [15], TransUNet combines the strengths of both Transformers and U-Net. The Transformer component encodes the tokenized image as an input sequence to extract global contexts, while the U-Net architecture is used to capture low-level visual cues, effectively preserving fine spatial details. Motivated by the Swin Transformer’s [16] success, Swin-Unet is introduced by to leverage the power of Transformer, based on a symmetric Encoder-Decoder architecture with skip connections.

However, almost all the above works have focused on improving the performance of the network, but they consume more computing resources and require more parameters, leading to higher memory usage, which poses challenges in resource-constrained medical environments. In this work, we focus on addressing this issue by designing an efficient network with reduced computational overhead, fewer parameters, and lower GFLOPs while also maintaining a good performance, by discarding a significant number of the Fourier coefficients from high frequencies.

### B. Fourier Frequency Learning

Frequency learning has great potential due to its brightness insensitivity and displacement invariance. The DC component in the spectrum represents the average brightness of the image, breaking the correlation between image content and brightness, and prompting the model to focus more on the target shape and structure. The Fast Fourier Transform (FFT) is a fast algorithm for computing the discrete Fourier transform of a sequence. Recently, several studies have applied the Fast Fourier Transform to images, utilizing frequency information to enhance both model performance and efficiency. Rao et al. propsed the Global Filter Network (GFNet), which learns long-term spatial dependencies in the frequency domain with log-linear complexity. They utilized FFT as the alternative to self-attention modules in the vision transformers [1]. Deep Fourier Up-Sampling, a novel method proposed by [19], enables the integration of features at different resolutions in the Fourier domain and addresses the challenges of up-sampling in the Fourier domain. In [20], GLFNET replaces self-attention mechanism with a combination of global-local filter blocks to optimize model efficiency. Some of these efforts based on well-known Convolution Theorem states that circular convolutions in the spatial domain are equivalent to pointwise products in the Fourier domain, to develop fast algorithms for training and inference. Mathieu et al. [22] present a simple algorithm to efficiently perform all convolutions as pairwise products by computing the Fourier transforms of the matrices in each set once. Inspired by [22], Adam et al. [23] proposed band-limited training, which not only effectively control the resource usage (GPU and memory) but also retain high prediction accuracy, through discarding high frequency spectrum. Nevertheless, most prior works encounter several issues. They have not fully utilized and tested all the advantages of the Fourier Frequency Domain, particularly in noise reduction. Noise in medical images is commonly caused by the malfunction of the sensor and other hardware in the process of forming, storing or transmitting images [25]. Noise causes degradation in image quality, which can affect the accuracy of medical diagnoses. Our proposed model, LightMed, is evaluated using advanced datasets with added noise in the images and has outperformed other models in nearly all metrics.

### C. Adversarial Attacks

Despite the significant contributions of deep neural networks across various AI fields, they remain vulnerable to carefully-engineered adversarial examples (attacks) with small imperceptible perturbations [39], [51], [52], [54]. This hazard has quickly become one of the most prominent areas of research in deep learning security community. Adversarial attacks exist across numerous data types, including images, audio, text, and other inputs [51], [52], [55]. These malicious examples can be generated in either a white-box setting [51], [56], [57], where an attacker has full access to the network model’s parameters, or a black-box setting [58], [59], where access is restricted strictly to the network’s inputs and outputs. Recent studies have confirmed that medical deep learning systems can also be compromised by adversarial attacks [52], [54]. Ma et al.[54] conducted experiments showing that adversarial attacks in the medical domain can succeed more easily than those on natural images due to complex biological textures and the overparameterization of state-of-the-art deep neural networks. Moreover, medical systems are notoriously difficult to update, making it challenging to implement new defenses [52]. Considering the huge healthcare economy, numerous defense models have been developed in medical image domain. Ren et al. [60] adopt adversarial defense to augment the training data, increase the robustness of the network for Brain MR Image Segmentation task. Likewise, [61], [63] mainly focus on adversarial training for improving intrinsic network robustness. Another approach attempts to distinguishing adversarial examples from normal clean examples in advance of being passed into model [54], [64], [65]. Some studies have shown the ancillary benefits of compression and distillation in resisting adversarial attacks [66]–[68]. These prior works suggest that adversarial attacks on neural networks often involve high-frequency perturbations of input data. Driven by these insights, our experiments demonstrate that LightMed, which directly learns in the frequency domain and harnesses low-frequency values, has robust capabilities in defending against adversarial examples.

#### a) Fast Gradient Sign Method (FGSM)

The FGSM algorithm [45] generates adversarial examples by adding a scaled sign of the gradient of the loss function with respect to the input, computed at the original data point *z*_0_. This perturbation is applied in a single step of magnitude *ε*:

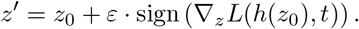

#### b) Basic Iterative Method (BIM)

BIM [47] extends FGSM by applying multiple smaller perturbations. Instead of a single large step, BIM refines the adversarial example through iterative updates using a step size *η*:

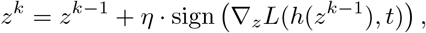

where *z*^*k*^ represents the adversarial example at the *k*-th iteration, starting from *z*^0^ = *z*_0_. The step size *η* is typically chosen in the range *ε/K* ≤ *η* ≤ *ε*, with *K* being the total number of iterations.

### c) Projected Gradient Descent (PGD)

PGD [46] improves upon BIM by adding a projection step to ensure the perturbations stay within allowed bounds. After each iteration, the adversarial example is projected back onto the *ε*-ball around the original input *z*_0_:

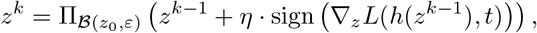

where 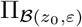 denotes the projection operator onto the ball of radius *ε* around *z*_0_. The initial point is often set as *z*^0^ = *z*_0_ + *U* (−*ε, ε*), where *U* (−*ε, ε*) is the uniform distribution in the range [−*ε, ε*]. PGD is considered one of the most effective first-order attack methods.

### D) Complex Architecture

Complex networks can be divided into two categories. First, there are networks where the input is complex and the model operates on real numbers. These networks convert complex numbers into real form before processing them, by stacking the real and imaginary parts into separate channels. In [22], [23], the convolution function is expedited by transforming the filter and the input into the frequency domain, followed by element-wise multiplication of both frequency spectra. On the other hand, there are networks where the input is real and the model operates on complex numbers. Trabelsi et al. [24] present a general formulation for the building components of complex-valued deep neural networks and apply it to the context of feed-forward convolutional networks and convolutional LSTMs. Their model relies on complex convolutions, complex batch-normalization and complex weight initialization strategies. LightMed is capable of handling complex inputs through complex operations and directly learning complex-valued spectra using complex computing networks, thus optimizing the use of complex spectral information.

## III. Preliminary

### A. Fourier Transform

We first give a brief review of the 2D Discrete Fourier Transform (DFT) that serves as an prevalent technique for investigating or modifying a signal. Fourier transform is based on the idea that functions of suitable properties can be represented with a linear combinations of trigonometric functions [36]. DFT of 2-dimensional signal *f* ∈ R*M ×N* is defined as:

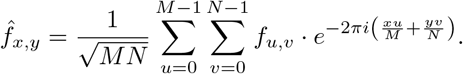

Correspondingly, the 2D Inverse DFT (2D-IDFT) is formulated as:

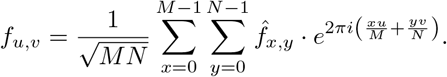

Both DFT and IDFT can be accelerated with their fast version, Fast Fourier Transform (FFT) algorithm [38] by James Cooley and John Tukey in 1965. FFT technique is widely applied in various fields, including JPEG data compression, solving partial differential equations, noise suppression.

### B. Band-limiting technique

Our model, LightMed, operates directly in the Fourier domain, taking transformed frequency data as its input. However, preserving the original frequency domain representation would pose challenges in computation and resource usage, as each element consists of both real and imaginary components. To overcome these challenges, we reduce the dimensionality of the inputs through compression while preserving sufficient image quality based on the following two properties of Fourier Transforms:

- **Conjugate Symmetry Property:** One of the key attributes of the Fourier Transform is the Conjugate Symmetry property. As a result, when we have the left half of our frequency map, the reduced number of degrees of freedom enables us to reconstruct the right half. This effectively allows us to store approximately half the parameters that would otherwise be required.
- **Low-pass Filter:** Since the Fourier coefficients of smooth background functions decay, most of their information is concentrated in the low-frequency components [36].

A common method for image compression involves converting an image to the frequency domain and applying a low-pass filter to suppress the high-frequency components of its Fourier transformation. By discarding enough of these high-frequency components, we reduce the amount of data needed to describe the image’s frequency content.

When the DC component is located in the top-left corner, these low frequencies are typically found at the four corners of the spectrum. Building on these approaches, we compress the inputs by discarding the right half of the frequency map, leveraging the Conjugate Symmetry Property, while retaining the low-frequency values located in the top-left and bottom-left corners, as depicted in Figure 1. While this reduces computational costs and memory usage, discarding too many high-frequency components can degrade image quality.

**Fig 1.**
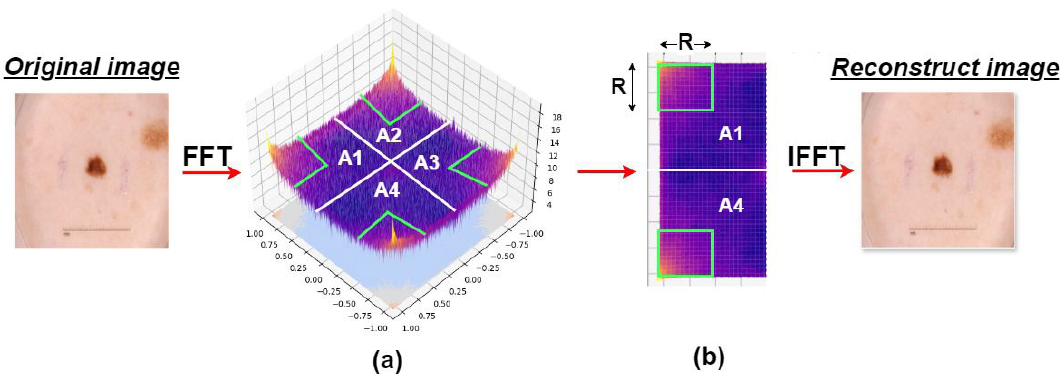
Overall image compression. **(a)** The log-amplitude map of the Fourier frequency domain is shown. Notably, the low-frequency values are predominantly concentrated at the four corners of the spectrum. **(b)** To optimize data storage, only the frequencies at two of the corners are retained. After applying the inverse 2D Fourier transform, the reconstructed image maintains sufficient fidelity to the original, despite the reduced data.

## IV. Method

### A. Overall Architecture

Our network architecture is illustrated in Fig.2. Our model features a U-shaped architecture similar to UNet. LightMed enables us to extract features directly from the frequency domain. Initially, the input image data is in the pixel domain, but after processing through a low-pass filter layer that employs the fast Fourier transform, we obtain a textured frequency space image map. The dimensions are reduced from *B ×* 3 *× H × W* to *B ×* 3 *×* 2*R × R*, where R is smaller than both H and W, as illustrated in Fig.2. The left side of our network, the Encoder, comprises five Complex Conblock layers. These layers reduce image dimensions in steps 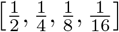 through four Anti-Aliasing downsampling transformations [62]. These transformations are crucial for retaining effective weights during the learning process, and simultaneously, the number of filters incrementally increases. At the crucial bottleneck at the lower right, we introduce a novel block, CCBAM, which enhances the architecture’s ability to learn both spatial and channel features in the frequency domain. The Decoder, located on the right side, consists of four Conv-Blocks that up-sample the data. Ultimately, the output is a prediction mask in the frequency domain. However, to visualize the mask in the pixel domain, we pad the empty positions in the frequency domain and then convert it back to the desired output format in the frequency domain.

**Fig 2.**
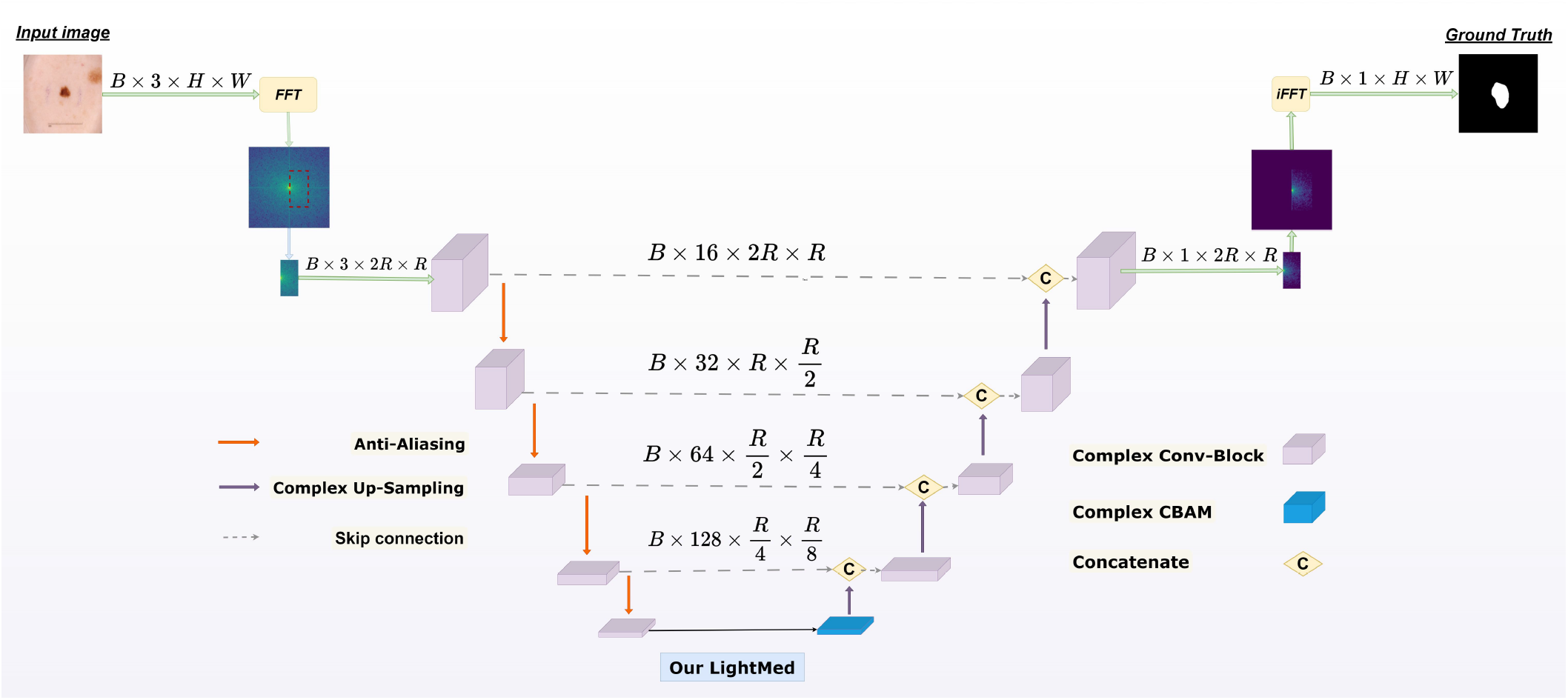
Our overall LightMed architecture learning on Frequency Domain.

### B. Complex Conv-Block

The Complex Conv-block, inspired by the Conv-Block proposed by Ronneberger et al. [3] and illustrated in Figure 3, consists of two layers of Complex Conv2d. The kernels in these layers sweep over both the real and imaginary domains of the image’s frequency space to produce corresponding real and imaginary feature maps. These convolutional layers are followed by activation functions such as ReLU and often include additional batch normalization layers.

**Fig 3.**
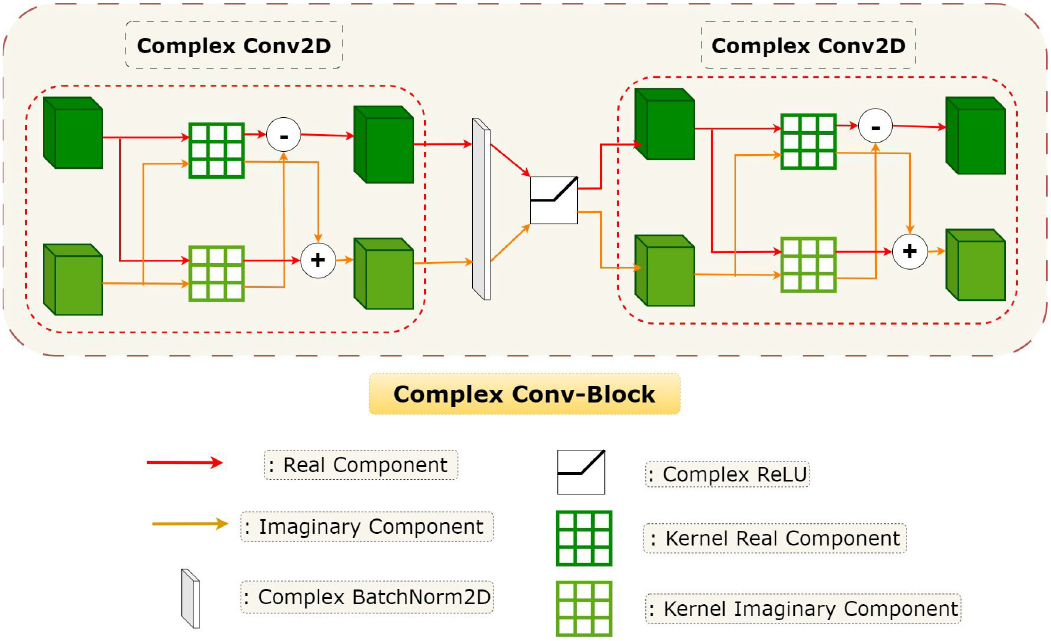
Our ComplexConv-Block.

Specifically, the use of ComplexReLU [24] helps prevent the vanishing gradient problem during training. Furthermore, ComplexBatchNorm2d [24] enhances the stability and accelerates the learning speed of the neural network by reducing the internal covariate shift in the frequency domain.

#### a. Complex Conv2D

In the spectrogram, the complex matrix *P* = *a* + *bi* consists of a real component *a* and an imaginary component *b*. These components are input into the network to perform an operation analogous to real-valued 2D convolution but within the complex domain. Simultaneously, two sets of convolution kernels *c* and *d* are introduced to represent the real and imaginary parts of the complex convolution kernel *Q* = *c* + *di*. The complex convolution operation can be expressed as:

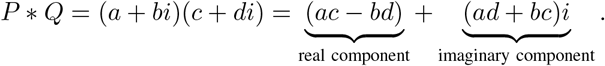

#### C. Complex Convolutional Block Attention Module

When processing an input feature map *F* in the frequency domain (where *F* ∈ ℂ^*C×H×W*^), we decompose *F* into its real component *F*_*r*_ and imaginary component *F*_*i*_ (both *F*_*r*_, *F*_*i*_ ∈ ℝ^*C×H×W*^). These components are then passed through attention mechanisms.

The Complex Convolutional Block Attention Module (CCBAM) employs a two-step complex attention mechanism. Initially, it derives a 1D complex channel attention map *M*_*C*_ ∈ ℂ^*C×*1*×*1^ (with real and imaginary parts *M*_*Cr*_, *M*_*Ci*_ ∈ ℝ^*C×*1*×*1^). Subsequently, it computes a 2D complex spatial attention map *M*_*S*_ ∈ ℂ^1*×H×W*^ (where *M*_*Sr*_, *M*_*Si*_ ∈ ℝ^1*×H×W*^), as depicted in Fig.4.

The attention processing stages can be concisely represented by the following equations:

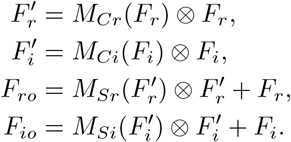

Here, ⊗ denotes element-wise multiplication. This operation involves appropriate broadcasting: channel attention is expanded across spatial dimensions, and spatial attention is expanded across channel dimensions. The resulting *F*_*ro*_ and *F*_*io*_ represent the finely adjusted outputs. Figures 4b) and 4c) illustrate the computational workflow for generating each type of attention map. Further details regarding each attention module are discussed subsequently.

**Fig 4.**
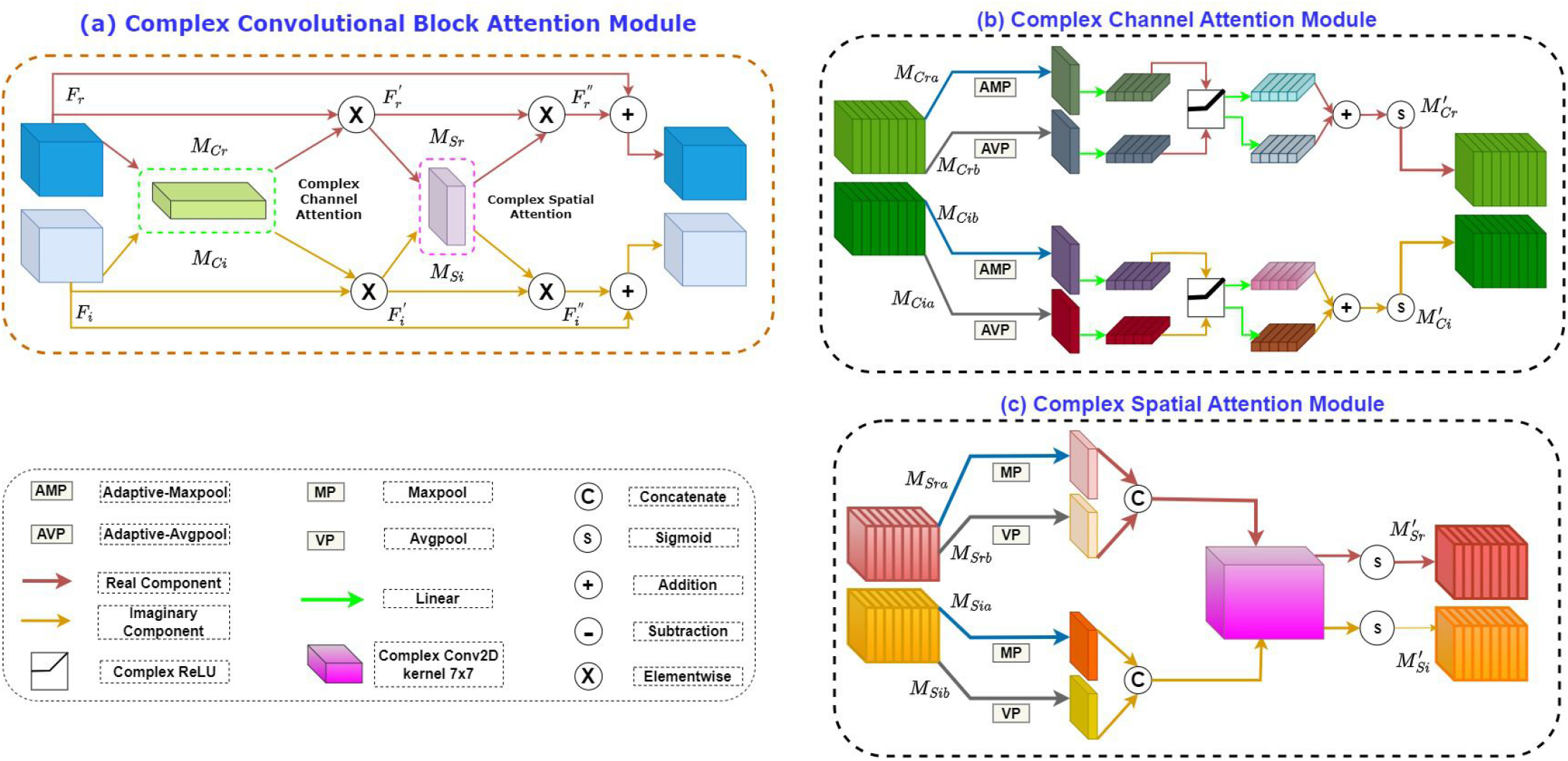
Our Complex Convolutional Block Attention Module, Complex Channel Attention Module and Complex Spatial Attention Module

#### a. Complex Channel Attention Module

Building upon the Channel Attention Module mechanism [2], our Complex Channel Attention Module generates a complex attention map by leveraging the inter-channel relationships of both real and imaginary features. In this approach, each channel of the feature map is divided into real and imaginary components. The channel attention mechanism focuses on identifying the significant features when the input image is transformed into the frequency domain. To compute the channel attention efficiently, we first reduce the spatial dimensions of the input feature map using the average pooling method. This technique helps in effectively capturing the extent of the target object. Additionally, prior research has shown that max pooling provides critical insights by gathering distinctive features across pixel layers in neural networks. This is particularly beneficial for enhancing channel-wise attention, even when operating in the frequency domain. Therefore, our method concurrently employs both average pooling and max pooling for the real and imaginary channels within the frequency domain. This dual approach ensures a more comprehensive and nuanced feature extraction, improving the overall representational power of the network. Detailed operational steps are outlined below.

Initially, we collect spatial information from the feature map through both average-pooling and max-pooling operations, resulting in two distinct spatial context descriptors: AVG (average-pooled features) and AMP (max-pooled features). These descriptors are then passed through a shared network to generate our channel attention map (*M*_*Cr*_, *M*_*Ci*_) ∈ R^*C×*1*×*1^. The shared network comprises a multi-layer perceptron (MLP) with a single hidden layer. To minimize parameter overhead, the hidden layer’s activation size is reduced to R^*C/r×*1*×*1^, where *r* represents the reduction ratio. After processing each descriptor through the shared network, the resulting feature vectors are combined using element-wise summation. In short, the complex channel attention is computed as:

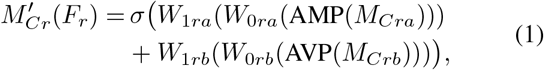

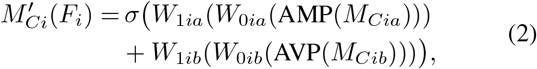

where (*W*_0*ra*_, *W*_0*rb*_, *W*_0*ia*_, *W*_0*ib*_) ∈ ℝ^*C/r×C*^, (*W*_1*ra*_, *W*_1*rb*_, *W*_1*ia*_, *W*_1*ib*_) ∈ ℝ^*C×C/r*^ and *σ* denotes the sigmoid function. Note that the MLP weights, (*W*_0*ra*_, *W*_0*rb*_, *W*_0*ia*_, *W*_0*ib*_) and (*W*_1*ra*_, *W*_1*rb*_, *W*_1*ia*_, *W*_1*ib*_), are shared for both inputs and the ComplexReLU activation function is followed by (*W*_0*ra*_, *W*_0*rb*_, *W*_0*ia*_, *W*_0*ib*_).

#### b. Complex Spatial Attention Module

Inspired by the Spatial Attention Module [2], we construct the Complex Spatial Attention Module (CSAM) which is a complex spatial attention map in the frequency domain by utilizing the spatial relationship between features. Different from complex channel attention, complex spatial attention focuses on ‘where’ is the information part of the real-to-virtual feature map, complementing complex channel attention. To compute complex spatial attention, we first apply average pooling and max pooling operations along the channel axis and concatenate them to generate an effective feature descriptor. On the concatenated feature representation, we apply a convolution operation using a specific layer to generate a spatial attention map with 2 real and imaginary domains (*M*_*Sr*_(*F*), *M*_*Si*_(*F*)) ∈ ℝ^*H×W*^ encoding where to emphasize or suppress. We describe the operation in detail below. We aggregate the channel information of the feature map using two pooling operations, generating two 2D maps over both the real and imaginary . domains in frequency space: 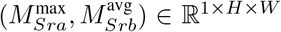 and 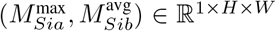Each map represents the average pooled features (VP) and the maximum pooled features (MP) over the entire channel. They are then concatenated and convolved by a standard convolutional layer, creating our 2D spatial attention map. In summary, spatial attention is computed as follows:

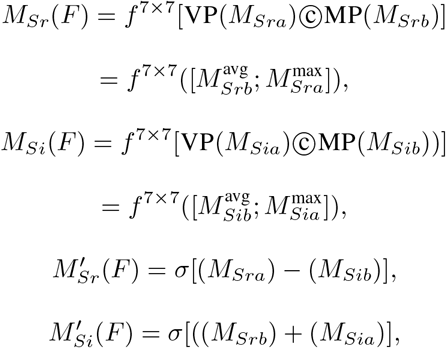

where *σ* denotes the sigmoid function, ?c denotes the concatenate and *f* ^7×7^ represents a convolution operation with the filter size of 7 *×* 7.

## V. Experiments

### A. Datasets

To evaluate the effectiveness of LightMed, we test it on four publicly available medical image datasets. In the deep learning, improve an optimal the segmentation of the model, we also albumentation training dataset: Horizontal Flip, Vertical Flip, Random Rotate 90, and Shift Scale Rotate. All image and mask sizes are both resized to 256 *×* 256. We propose a state of the art benchmark for five datasets based on additional Gaussian for testing dataset. Finally, we conduct additional experiments on the ISIC-2018 dataset.

#### 1) ISIC 2018 dataset

ISIC 2018 [31] provides a separate data set for the task of skin disease edge region segmentation for the first time. The ISIC 2018 dataset has become the main benchmark for evaluating skin lesion image segmentation algorithms, which contains 2594 RGB skin lesion images. The original training set in this study is split into a new training set, validation set and test set with 70%, 10% and 20%.

#### 2) ISIC 2017 dataset

In order to evaluate the effectiveness of LightMed on the skin cancer image segmentation task, we selected the ISIC-2017 dataset, which contains 2000 training images, 150 validation images, and 600 test images [32]. Ground truth and patient metadata are provided for both training and test sets, indicating if the lesion is one of four class groups: (1) melanoma; (2) nevus or seborrheic keratosis; (3) seborrheic keratosis; or (4) melanoma or nevus. The patient’s approximate age and gender are also provided as additional metadata [53].

#### 3) PH2 dataset

The PH2 dataset [50] comprises 200 dermoscopic images in 8-bit RGB format, with resolutions ranging from 553 × 763 to 577 × 769 pixels. We randomly assigned 140 images for training purposes, 20 image for validation and the remaining 40 images for testing.

#### 4) Covid-19 dataset

COVID-19 database [40]–[43], comprising 21,165 chest X-ray images with lung masks for segmentation. Considering the computational cost, we sample 2,750 Xray images and corresponding lung masks from the COVID19 database for medical segmentation [49]. We randomly assigned 70% images for training purposes, 10% image for validation and the remaining 20% images for testing.

#### 5) ACDC dataset

The Automated Cardiac Diagnosis Challenge (ACDC) [48] dataset encompass a series of short-axis sections, covering the range from the base to the apex of the left ventricle, with slice thickness ranging from 5 to 8 millimeters. the ACDC dataset includes 100 samples, with 70 for training, 10 for validation, and 20 for testing. However, we have taken steps to increase the effective size of the dataset by slicing the 3D volumes along the z-axis, resulting in a larger number of 2D slices. Specifically, the 70 training samples of the ACDC were converted into 1,312 slices for training, and the 10 validation samples were split into 202 slices for validation, and the 20 test samples were transformed into 388 slices for testing.

### B. Evaluation Metrics

Mean Intersection over Union (mIoU) and F1-score are standard metrics commonly used in medical image segmentation. mIoU is particularly prevalent in competitions for comparing the performance of different models. To provide a more exhaustive comparison between the performance of LightMed and other popular models, we calculate each of these metrics in our experiment.

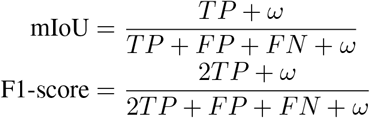

Here, *TP*, *FP*, *FN*, and *TN* represent true positive, false positive, false negative, and true negative, respectively. Note that *ω* is set to 1 *×* 10^−6^ to avoid division by zero.

### C. Loss function and Comparison settings

#### 1) Loss function

To ensure a fair comparison, we reran all experiments using a combined loss function that integrates the Binary Cross-Entropy (BCE) loss with the Dice loss, assigning equal weights *α* = *β* = 1:

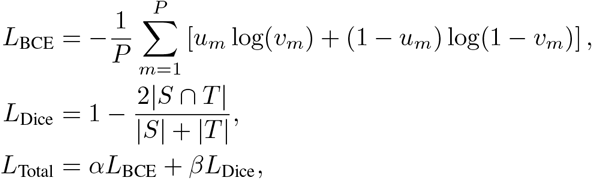

where *P* is the total number of samples, *u*_*m*_ represents the true labels, and *v*_*m*_ denotes the predicted probabilities. The terms |*S*| and |*T*| correspond to the sizes of the ground truth and predicted segmentations, respectively.

#### 2) Training Process

LightMed was developed with PyTorch version 2.4.0 and employed NVIDIA TESLA P100 GPUs. We used CPU Intel Xeon 2.00Ghz of Kaggle testing Inference times (ms) model proposed. During training, the network was optimized using the AdamW algorithm with an initial learning rate of 1e-3 and a batch size of 16. In addition, the CosineAnnealingLR [44] is employed as the scheduler with a maximum number of iterations of 50 and a minimum learning rate of 1e-5.

### D. Comparative results

#### 1) Evaluation on the ISIC-2017 and ISIC-2018 Dataset

The experimental results presented in Table I demonstrate that our proposed LightMed model exhibits superior performance compared to CNNs and Transformer methods in the specific biomedical image segmentation task of skin lesion segmentation. On the ISIC-2017 dataset, LightMed achieves outstanding results with an F1 score of 80.38% and an IoU of 67.21%, outperforming the CNN-based U-Net [3], which has an F1 score of 80.08% and an IoU of 66.78%, as well as Transformer models like Trans-UNet [15] with an F1 score of 71.14% and an IoU of 58.67%.

**TABLE I.** Performance Comparison LightMed model on the ISIC-2017 and ISIC-2018 dataset.

| Method | Params(M)↓ | Memory Usage(MB)↓ | GFLOPs↓ | ISIC-2017 |  | ISIC-2018 |  |
| --- | --- | --- | --- | --- | --- | --- | --- |
|  |  |  |  | %F1 ↑ | %mIoU ↑ | %F1 ↑ | %mIoU ↑ |
| UNet [3] | 33.12 | 132.55 | 15.35 | 80.08 ± 0.613 | 66.78 ± 0.852 | 23.92 ± 12.93 | 14.04 ± 08.65 |
| UNet++ [4] | 9.04 | 36.26 | 81.67 | 75.82 ± 03.67 | 61.17 ± 04.66 | 59.06 ± 13.41 | 42.89 ± 13.02 |
| ResUNet [5] | 46.41 | 185.83 | 36.57 | 11.04 ± 06.27 | 06.37 ± 03.53 | 19.86 ± 19.12 | 15.45 ± 13.46 |
| Attention Unet [14] | 34.87 | 139.64 | 66.63 | 60.71 ± 13.62 | 44.61 ± 24.51 | 24.51 ± 16.19 | 14.80 ± 11.31 |
| TransUNet [15] | 67.07 | 268.46 | 32.63 | 71.14 ± 03.51 | 58.67 ± 03.67 | 37.70 ± 13.55 | 23.93 ± 10.58 |
| CaraNet [21] | 46.64 | 187.5 | 11.50 | 76.23 ± 04.07 | 61.72 ± 05.29 | 65.23 ± 12.79 | 49.41 ± 13.36 |
| PraNet [33] | 32.54 | 130.73 | 06.96 | 78.51 ± 02.43 | 64.68 ± 03.30 | 74.15 ± 02.79 | 58.98 ± 03.54 |
| LightMed +w/o CCBAM | 3.98 | 16.01 | 1.72* | 80.14 ± 1.51 | 66.94 ± 1.82 | 81.35 ± 0.58 | 68.94 ± 0.64 |
| **LightMed (Ours)** | **3.98** | **16.01** | **1.72** | **80.38 ± 1.14** | **67.21 ± 1.58** | **81.75 ± 0.43** | **69.41 ± 0.61** |

\*\*: GFLOPs of CCBAM accounts for $8 \times 10^{-4}\%$ of LightMed

Additionally, on the ISIC-2018 dataset, LightMed delivers impressive results with an F1 score of 81.75%, significantly better than CNN methods such as Attention U-Net [14] with an F1 score of 24.51% and pretrained methods like CaraNet with an F1 score of 65.23%. LightMed is considered a lightweight model with only 3.98 million parameters, compared to larger models like U-Net (33.12M), Attention U-Net (34.87M), and Transformer models with large pretrained parameters such as TransUNet [15] (67.07M) and CaraNet [21] (46.64M).

Furthermore, our model exhibits significantly smaller storage requirements of just 16.01 MB and computational needs of only 1.72 GFlops, which are minimal compared to other methods listed in Table I. In our proposal, this result table employs Gaussian noise level 4 for the ISIC-2017 dataset and noise level 1 for the ISIC-2018 dataset. Additionally, we show the performance of binary skin lesion segmentation on the ISIC-2017 and ISIC-2018 datasets in Figure 5.

**Fig 5.**
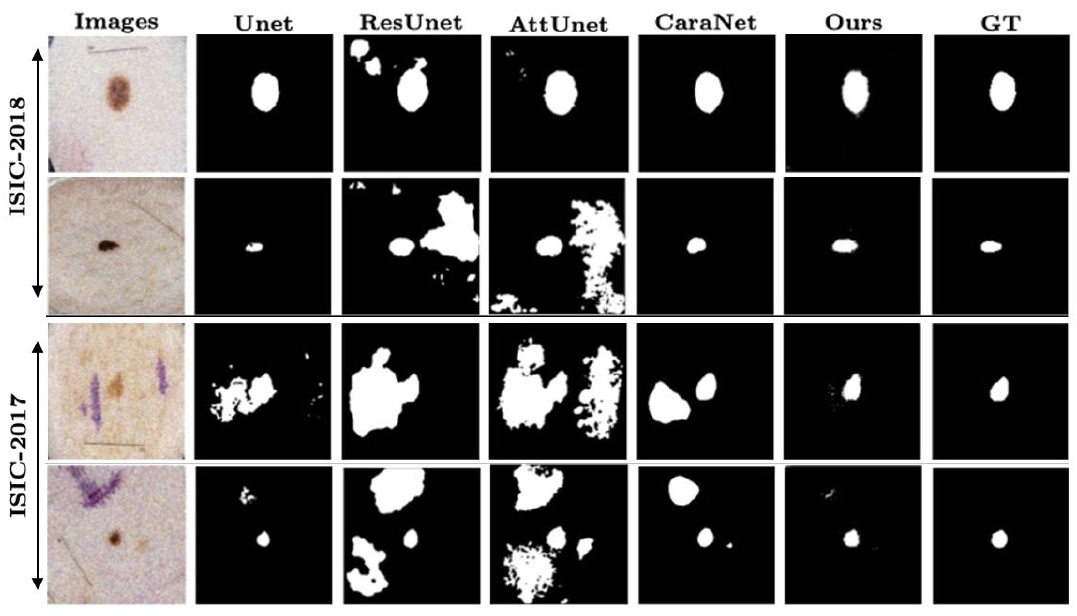
Qualitative results LightMed with other methods on ISIC-2017 and ISIC-2018 dataset.

**Fig 6.**
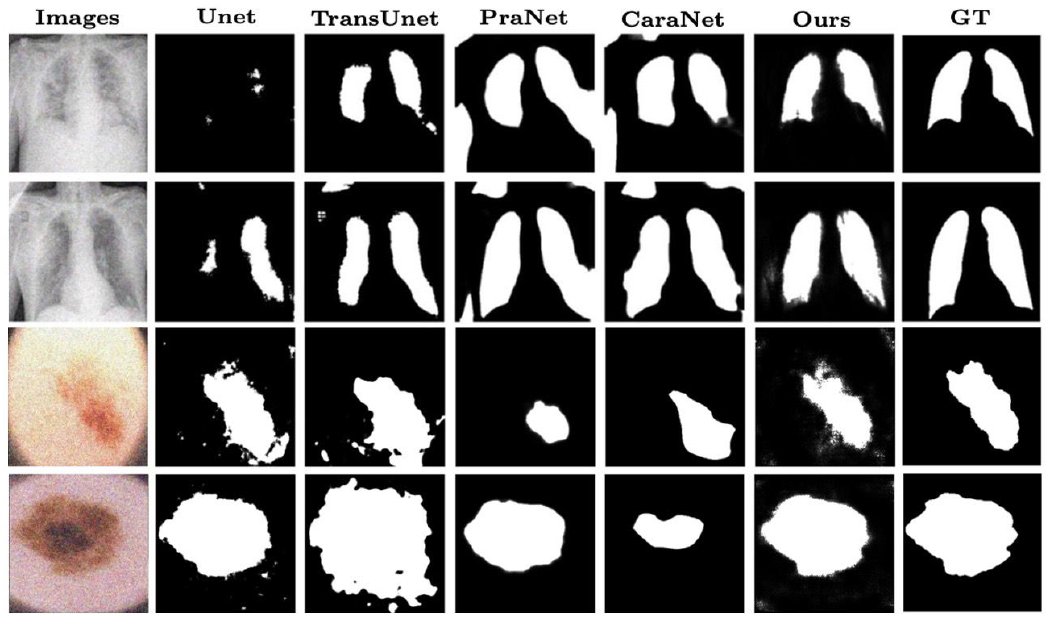
Qualitative results LightMed with other methods on Lung Covid-19 and PH2 dataset.

**Fig 7.**
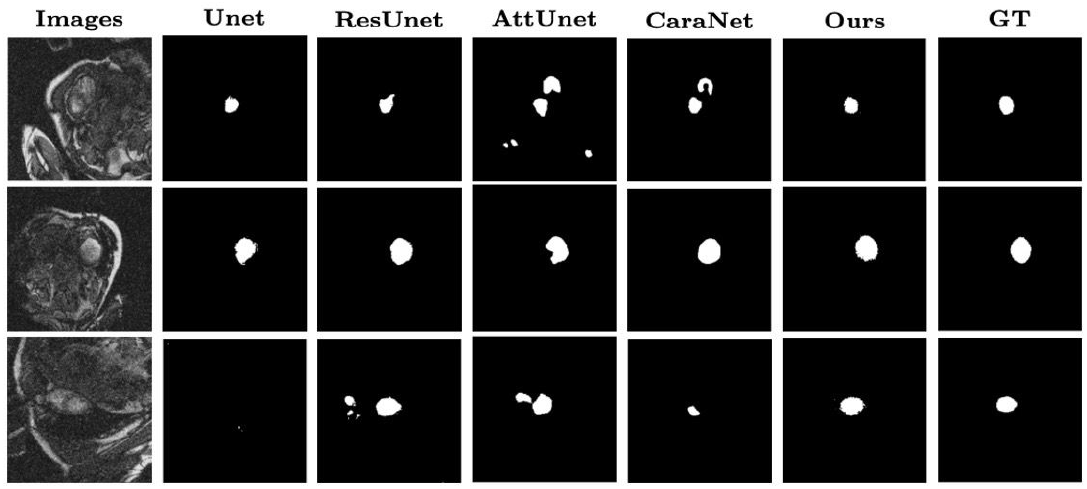
Qualitative results LightMed with other methods on the ACDC dataset.

#### 2) Evaluation on the Covid-19 and PH2 Dataset

In this benchmark, we evaluate the performance of our proposed LightMed model against other models using the PH2 dataset, which contains fewer samples compared to the skin lesion segmentation datasets discussed earlier. Additionally, we assess our architecture on a chest X-ray dataset for lung segmentation, as detailed in Table II. Notably, LightMed requires fewer parameters, less memory, and lower Gflops than the competing models. The experimental results presented in Table II demonstrate that LightMed outperforms other methods in both the Lung Covid-19 and PH2 datasets. On the Lung Covid-19 dataset, LightMed achieves an impressive F1 score of 93.02 *±* 0.05% and an IoU of 86.94 *±* 0.08%, significantly surpassing models such as U-Net, which has an F1 score of 35.52 *±* 20.78% and an IoU of 23.21 *±* 15.98%, and TransUNet [15], which records an F1 score of 79.10 *±* 4.38% and an IoU of 65.59 *±* 6.01%. These results highlight the superior performance and efficiency of LightMed in biomedical image segmentation tasks.

**TABLE II.** Performance Comparison of LightMed model on the Covid-19, PH2, and ACDC (LV) datasets.

| Method | Params(M) ↓ | Memory Usage(MB) ↓ | GFLOPs ↓ | Lung Covid-19 |  | PH2 |  | ACDC (LV) |  |
| --- | --- | --- | --- | --- | --- | --- | --- | --- | --- |
|  |  |  |  | %F1 ↑ | %mIoU ↑ | %F1 ↑ | %mIoU ↑ | %F1 ↑ | %mIoU ↑ |
| UNet [3] | 33.12 | 132.55 | 15.35 | 35.52 ± 20.78 | 23.21 ± 15.98 | 79.65 ± 01.81 | 66.23 ± 02.50 | 21.00 ± 19.41 | 15.09 ± 14.41 |
| UNet++ [4] | 09.04 | 36.26 | 81.67 | 44.70 ± 24.28 | 31.03 ± 18.01 | 73.69 ± 08.09 | 58.94 ± 09.39 | 16.25 ± 13.86 | 9.36 ± 8.29 |
| ResUNet [5] | 46.41 | 185.83 | 36.57 | 39.37 ± 35.28 | 29.25 ± 27.07 | 72.52 ± 09.56 | 57.69 ± 10.71 | 27.57 ± 12.92 | 13.18 ± 8.46 |
| Attention Unet [14] | 34.87 | 139.64 | 66.63 | 50.64 ± 13.15 | 34.73 ± 11.76 | 72.26 ± 08.00 | 57.17 ± 09.59 | 21.19 ± 16.88 | 12.65 ± 10.63 |
| TransUNet [15] | 67.07 | 268.46 | 32.63 | 79.10 ± 04.38 | 65.59 ± 06.01 | 69.97 ± 03.42 | 53.92 ± 04.05 | 37.47 ± 16.13 | 24.02 ± 12.17 |
| CaraNet [21] | 46.64 | 187.5 | 11.50 | 71.22 ± 05.04 | 55.50 ± 06.17 | 63.73 ± 09.42 | 47.46 ± 10.02 | 56.05 ± 21.19 | 41.10 ± 19.69 |
| PraNet [33] | 32.54 | 130.73 | 06.96 | 66.24 ± 04.65 | 49.67 ± 05.25 | 76.73 ± 04.39 | 62.46 ± 05.93 | 55.80 ± 8.07 | 39.02 ± 7.72 |
| LightMed +w/o CCBAM | 3.98 | 16.01 | 1.72* | 92.99 ± 0.078 | 86.64 ± 0.082 | **88.60 ± 02.42** | **79.21 ± 03.97** | 78.13 ± 0.66 | 63.01 ± 0.89 |
| **LightMed (Ours)** | **3.98** | **16.01** | **1.72** | **93.02 ± 0.054** | **86.94 ± 0.081** | 88.20 ± 02.38 | 78.98 ± 03.85 | **77.90 ± 0.64** | **63.79 ± 0.86** |

On the PH2 dataset, LightMed also delivers impressive results with an F1 score of 88.20 *±* 2.38% and IoU of 78.98 *±* 3.85%, which is notably better compared to other methods. The remarkably low standard deviations of LightMed’s results, particularly on the Lung Covid-19 dataset, underscore the model’s exceptional stability and reliability across different runs. In contrast, the higher standard deviations of other models indicate greater performance variability, potentially leading to less reliable outcomes in some instances.

#### 3) Evaluation on the Left Ventrigle Segmentation on ACDC Dataset

Table II presents a comparative analysis of various models, including our proposed “LightMed,” on the LV segmentation task using the ACDC dataset. LightMed (Ours) is the most resource-efficient model, featuring the fewest parameters (3.98M) and the lowest memory usage (16 MB). In contrast, models such as TransUNet and PraNet have significantly higher parameter counts and memory requirements, with TransUNet utilizing 268.6 MB. Additionally, LightMed excels in computational efficiency, requiring only 1.72 GFLOPS. Conversely, ResUNet and Attention U-Net demand substantially more computational resources, with GFLOPS values of 36.57 and 66.63, respectively. Despite its lightweight architecture, LightMed achieves the highest performance metrics, recording an F1 score of 77.90 *±* 0.64% and an IoU of 63.79 *±* 0.86%, thereby outperforming all other models listed in Table II. Furthermore, TransUNet and UNet++ exhibit lower F1 and IoU scores, indicating reduced segmentation accuracy compared to LightMed.

**TABLE III.** Gaussian Noise Settings for Different Levels.

|  | Gaussian Noise setting increase level |  |  |  |  |
| --- | --- | --- | --- | --- | --- |
|  | Level 0 | Level 1 | Level 2 | Level 3 | Level 4 |
| Range for noise | - | (0.01, 0.015) | (0.015, 0.02) | (0.02, 0.025) | (0.025, 0.03) |
| Mean | - | 0 | 0 | 0 | 0 |
| Add noise per channel | - | TRUE | TRUE | TRUE | TRUE |
| Scaling factor for noise generation | - | 1.0 | 1.0 | 1.0 | 1.0 |
| Probability of applying the transform | - | 1.0 | 1.0 | 1.0 | 1.0 |

### E. Ablation Study

#### 1) Model stability when increasing noise

In this section, we will analyze the stability of the proposed model with increasing levels of Gaussian Noise added to the test set on the dataset. The settings for Gaussian Noise are described in Table Note that at level 0 we have not set Gaussian Noise, we use “-” instead. From level 1 to level 4 we gradually increase the Variance range for noise. With the mean of the default noise equal to 0.

At the same time, set true to add noise to each channel in the RGB image. Finally, Probability of applying the transform to ensure that every image in testing has noise. In this Fig. 8 illustrates the noise resilience of our method, showing consistent stability compared to other methods as the level of Gaussian noise increases.

**Fig 8.**
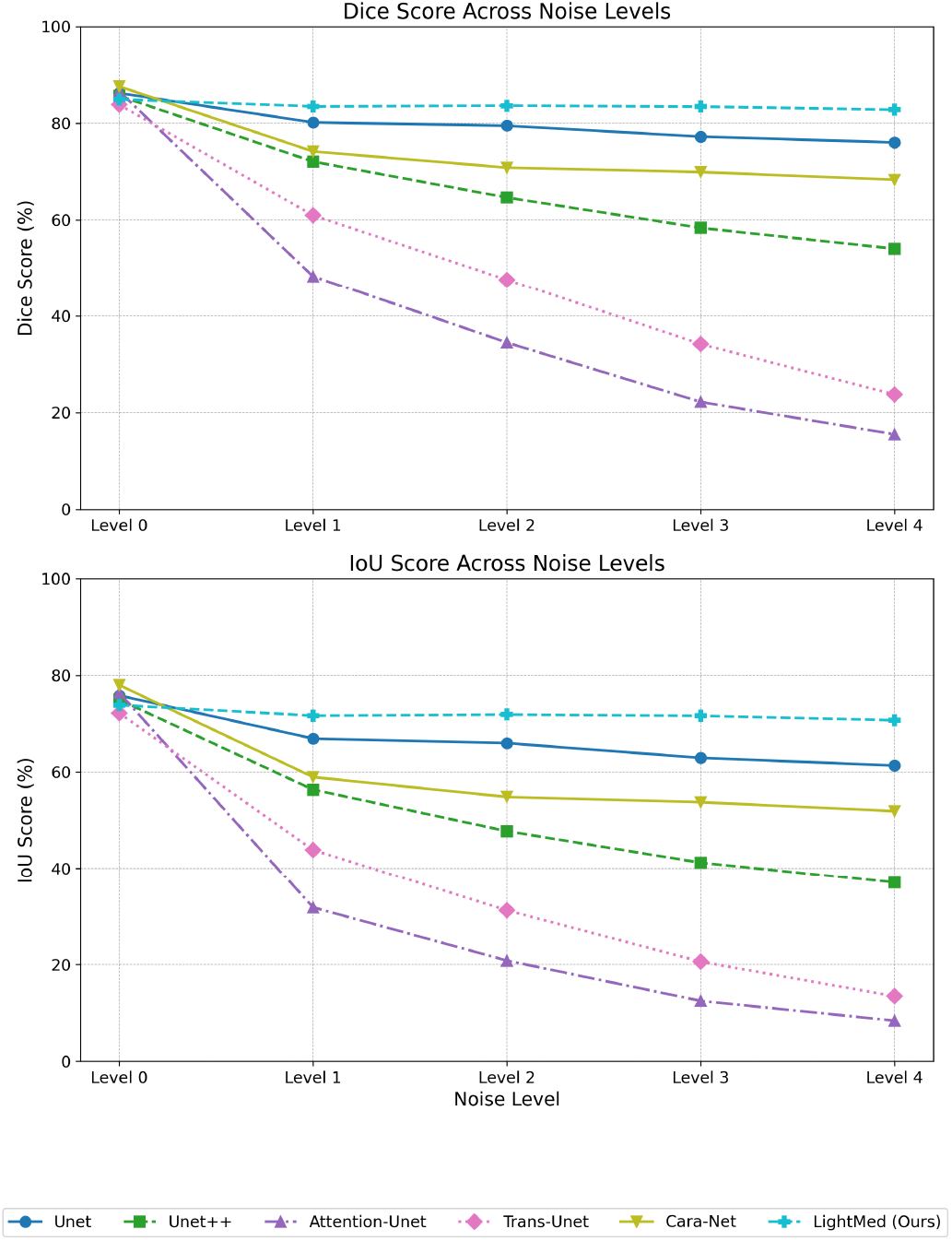
Qualitative results of different methods.

### 2) Analysis on the number of channels

**LightMed-L:** Featuring the most channels [32, 64, 128, 256, 512], LightMed-L contains the highest number of parameters (15.88M), the slowest inference speed (22.3 ms), and the greatest computational demand (6.84 GFLOPs). Despite these increases, it only slightly improves performance, attaining an F1 score of 83.87 and an mIoU of 72.21. **LightMed-S:** Utilizing the fewest channels [8, 16, 32, 64, 128], LightMed-S has the smallest number of parameters (0.99M), the fastest inference speed (22.7 ms), and the lowest computational cost (0.43 GFLOPs). Consequently, it achieves the lowest performance with an F1 score of 83.44 and an mIoU of 71.59. **LightMed (Ours):** Striking a balance with a moderate number of channels [16, 32, 64, 128, 256], LightMed incorporates 3.98M parameters and 1.73 GFLOPs, maintaining an inference speed of 22.6 ms. It delivers the highest performance, achieving an F1 score of 83.92 and an mIoU of 72.32, demonstrating an optimal trade-off between efficiency and accuracy.

#### 3) Analysis different R on LightMed

GFLOPS represents the computational cost of the model.Based on the Table V we can see lower GFLOPS indicates a more computationally efficient model. Inference Time ↓ refers to the time it takes for the model to make a prediction on a single input. Lower inference time is preferred, especially in real-time applications. GFLOPS when **R = 32** has the lowest computational cost (0.43 GFLOPS), making it the most efficient. Inference Time: **R = 32** is the fastest with 32.67 ms. F1 and mIoU: **R = 64** offers the best balance, with slightly higher F1 (81.75%) and mIoU (69.41%) scores without a significant increase in computational cost.

#### 4) Compare the performance stability of the architecture in noise environment combine Adversdial Attack

In this ablation study, we examine the robustness of the proposed LightMed model against other CNN and Transformer methods in the task of medical image segmentation. To enhance attack diversity, we introduce a Gaussian noise layer at level 1, as shown in Table 4, before applying the attack levels of FGSM, BIM, and PGD. This experiment highlights the model’s resilience not only to adversarial examples generated by FGSM, BIM, and PGD but also to other types of noise that may occur naturally, such as Gaussian noise.

**TABLE IV.** Analysis on the number of channels.

| Method | Channels | Params | Inf. Speed (in ms) ↓ | GFLOPs ↓ | %F1 ↑ | %mIoU ↑ |
| --- | --- | --- | --- | --- | --- | --- |
| LightMed-S | [8, 16, 32, 64, 128] | **0.99** | **38.15** | **0.43** | 80.23 | 66.98 |
| LightMed-L | [32, 64, 128, 256, 512] | 15.88 | 328.75 | 6.84 | 81.65 | 68.63 |
| **LightMed (Ours)** | [16, 32, 64, 128, 256] | 3.98 | 52.81 | 1.72 | **81.75 ± 0.43** | **69.41 ± 0.61** |

**TABLE V.** Comparison different R on LightMed.

| Method | GFLOPs ↓ | Inference times (ms) ↓ | %F1 ↑ | %mIoU ↑ |
| --- | --- | --- | --- | --- |
| LightMed [R = 32] | **0.43** | **32.67** | 81.37 | 68.60 |
| LightMed [R = 128] | 6.87 | 123.44 | 81.71 | 69.09 |
| **LightMed [R = 64] (Ours)** | 1.72 | 52.81 | **81.75 ± 0.43** | **69.41 ± 0.61** |

The attacks we consider include: 1) the single-step FGSM, 2) the iterative BIM, and 3) the strongest first-order attack PGD. Note that all these attacks are constrained by a predefined maximum perturbation *ϵ* according to the *L*_∞_ norm, meaning the perturbation on each input pixel does not exceed *ϵ* [54]. All three attacks are applied to the testing set with the added Gaussian noise at level 1. When an image is processed, the input gradient extractor feeds it into the pre-trained segmentation model to obtain the input gradients, which are then perturbed to maximize the network’s loss. The steps for FGSM, BIM, and PGD are progressively increased with *ϵ* = *x* * 0.1*/*255 where *x* = [2, 4, 6, 8].

As you can be seen in Table VI, with *x* = 2, LightMed still performs well in resisting Gaussian Noise + FGSM, achieving an F1 score of 75.02%, compared to CNN methods like UNet with an F1 score of 34.83%, or Transformers like TransUnet with an F1 score of only 29.68%. Tables VII and Tables VIII also demonstrate the superior performance of the LightMed model. Our model’s results remain unchanged when defending against the PGD method, which is considered stronger than BIM. The reason is that the amount of noise added to the initial image by PGD does not affect the LightMed model, as it operates in the frequency domain.

**TABLE VI.** Performance Comparison under FGSM attack + Gaussian Noise.

| Method | x = 2 |  | x = 4 |  | x = 6 |  | x = 8 |  |
| --- | --- | --- | --- | --- | --- | --- | --- | --- |
|  | %IoU ↑ | %Dice ↑ | %IoU ↑ | %Dice ↑ | %IoU ↑ | %Dice ↑ | %IoU ↑ | %Dice ↑ |
| Unet [3] | 14.14 | 24.78 | 13.24 | 23.38 | 12.42 | 22.09 | 11.69 | 20.92 |
| Unet++ [4] | 20.66 | 34.25 | 18.69 | 31.49 | 16.93 | 28.96 | 15.24 | 26.73 |
| ResUNet [5] | 21.08 | 34.83 | 18.89 | 31.78 | 17.21 | 29.36 | 11.69 | 27.43 |
| Attention Unet [14] | 07.61 | 13.75 | 07.19 | 13.43 | 06.82 | 12.78 | 06.48 | 12.11 |
| CaraNet [21] | 48.24 | 65.09 | 42.44 | 59.59 | 38.68 | 55.77 | 35.04 | 53.04 |
| PraNet [33] | 39.63 | 56.76 | 34.65 | 51.47 | 34.11 | 47.81 | 28.55 | 44.89 |
| TransUNet [15] | 17.42 | 29.68 | 15.69 | 27.12 | 14.23 | 24.91 | 12.99 | 23.00 |
| **LightMed (Ours)** | **60.02** | **75.02** | **56.15** | **71.92** | **53.15** | **69.41** | **50.69** | **67.29** |

**TABLE VII.** Performance Comparison under BIM attack + Gaussian Noise.

| Method | x = 2 |  | x = 4 |  | x = 6 |  | x = 8 |  |
| --- | --- | --- | --- | --- | --- | --- | --- | --- |
|  | %IoU ↑ | %Dice ↑ | %IoU ↑ | %Dice ↑ | %IoU ↑ | %Dice ↑ | %IoU ↑ | %Dice ↑ |
| Unet [3] | 12.35 | 21.99 | 07.79 | 14.46 | 03.66 | 07.06 | 01.77 | 03.48 |
| Unet++ [4] | 16.78 | 28.73 | 08.40 | 15.50 | 03.15 | 06.1 | 01.92 | 03.77 |
| ResUNet [5] | 16.77 | 28.72 | 09.58 | 17.48 | 05.10 | 09.71 | 04.14 | 0.795 |
| Attention Unet [14] | 06.81 | 12.75 | 04.65 | 08.89 | 02.05 | 04.01 | 00.97 | 01.92 |
| CaraNet [21] | 35.34 | 52.22 | 20.18 | 33.58 | 13.79 | 24.23 | 11.99 | 21.42 |
| PraNet [33] | 29.53 | 45.60 | 16.59 | 28.47 | 11.60 | 20.79 | 09.64 | 17.59 |
| TransUNet [15] | 14.05 | 24.63 | 07.09 | 13.24 | 02.05 | 04.02 | 00.54 | 01.08 |
| **LightMed (Ours)** | **51.98** | **68.41** | **36.33** | **53.29** | **25.09** | **40.12** | **20.38** | **33.85** |

**TABLE VIII.** Performance Comparison under PGD attack + Gaussian Noise.

| Method | x = 2 |  | x = 4 |  | x = 6 |  | x = 8 |  |
| --- | --- | --- | --- | --- | --- | --- | --- | --- |
|  | %IoU ↑ | %Dice ↑ | %IoU ↑ | %Dice ↑ | %IoU ↑ | %Dice ↑ | %IoU ↑ | %Dice ↑ |
| Unet [3] | 12.34 | 21.98 | 07.79 | 14.46 | 03.66 | 07.06 | 01.77 | 03.48 |
| Unet++ [4] | 16.77 | 28.73 | 08.41 | 15.51 | 03.15 | 06.11 | 01.92 | 03.76 |
| ResUNet [5] | 16.77 | 28.72 | 09.60 | 17.55 | 05.13 | 09.76 | 04.21 | 08.17 |
| Attention Unet [14] | 06.81 | 12.75 | 04.65 | 08.89 | 02.05 | 04.01 | 00.97 | 01.92 |
| CaraNet [21] | 35.34 | 52.23 | 20.18 | 33.59 | 13.78 | 24.23 | 11.99 | 21.42 |
| PraNet [33] | 29.53 | 45.60 | 16.59 | 28.47 | 11.60 | 20.79 | 09.64 | 17.59 |
| TransUNet [15] | 32.24 | 48.43 | 20.18 | 33.59 | 13.78 | 24.23 | 11.99 | 21.42 |
| **LightMed (Ours)** | **51.98** | **68.41** | **36.33** | **53.29** | **25.09** | **40.12** | **20.38** | **33.85** |

## VI. Future Works

A promising direction for extending the capabilities of LightMed is its integration with Large Language Models (LLMs) to build a Medical Vision-Language Model (VLM).

By combining LightMed’s segmentation results with VLMs, which merge visual information from medical images with textual medical knowledge, it becomes possible to create a comprehensive diagnostic tool. This integration would enable automated interpretation of segmented images, providing both visual outputs and accompanying diagnostic narratives. Leveraging transformer-based LLMs trained on medical data could further enhance this system, facilitating automated report generation and clinical decision support. This combination would significantly improve the tool’s utility in fields such as radiology and dermatology, where both images and text-based diagnoses play a crucial role.

## VII. Conclusion

In this paper, we present LightMed, a lightweight model in terms of parameters, gflops, inference times, and memory usage that is learned in the Fourier frequency domain. In addition, we present novel proposals for Complex Conv-Block convolutions and a Convolutional Block Attention Module that support frequency domain learning. Our study demonstrates that, first, for natural noise tasks such as Gaussian noise, it outperforms previous research models in the biomedical image segmentation problem through low-pass filter compression at the model input. Second, in additional experiments, the LightMed model also produces superior results even when encountering natural noise combined with Adversadial Attack.

**Dr. Truong Son Hy** is currently a tenure-track Assistant Professor in the Department of Computer Science, University of Alabama at Birmingham, United States. He has earned his PhD in Computer Science from University of Chicago, United States; and his BSc in Computer Science from E+ tv+ s Loránd University, Hungary. Prior to his current faculty position, he has worked as a lecturer and postdoctoral fellow in the Halicio+ lu Data Science Institute at University of California, San Diego; and a tenure-track Assistant Professor in the Department of Mathematics and Computer Science at Indiana State University, United States. His research focuses on graph neural networks, multiresolution matrix factorization, graph wavelets, deep generative models on graphs, group equivariant, and multiscale hierarchical models for scientific applications in the direction of **AI for Science and Engineering**.

**Viet Tien Pham** is currently a Research Assistant under the supervision of Dr. Truong Son Hy at the University of Alabama at Birmingham, United States. He earned his BSc degree in Automation Control Engineering from the School of Electrical and Electronic Engineering, Hanoi University of Science and Technology, Vietnam. His research focuses on foundational models for scientific applications, particularly in the field of **AI for Science and Engineering**.

**Minh Hieu Ha** is currently a student in the Talented Program of Computer Science at the School of Information and Communication Technology, Hanoi University of Science and Technology. He serves as a Research Assistant under the supervision of Dr. Truong Son Hy at the University of Alabama at Birmingham, United States. His research focuses on foundational models for scientific applications, with an emphasis on **AI for Science and Engineering**.

**Bao V. Q. Bui** is currently a student in the Honor Program at Ho Chi Minh City University of Technology (HCMUT), VNU-HCM. He is interning as a Research Assistant under the supervision of Dr. Truong Son Hy at the University of Alabama at Birmingham, United States. His research focuses on foundational models for scientific applications, with an emphasis on **Medical Imaging**.

## APPENDIX A Appendix I

### Implementation details

#### A. Fast Fourier Transforms

##### Fast Fourier Transform 1D

(Cooley and Tukey, 1965) was first invented by Carl Friedrich Gauss and then rediscovered by Cooley and Tukey (the radix-2 Decimation In Time or DIT case). DFT is defined by the formula:

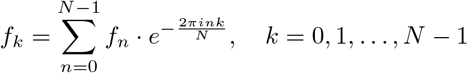

Radix-2 DIT first computes the DFTs of the even-indexed inputs (*f*_2*m*_ = *f*_0_, *f*_2_, …, *f*_*N*−2_) and of the odd-indexed inputs (*f*_2*m*+1_ = *f*_1_, *f*_3_, …, *f*_*N*−1_) and then combines those two results to produce the DFT of the whole sequence:

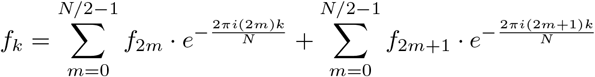

The equation can be represented as:

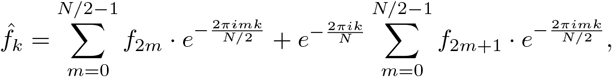

that can be written as

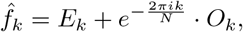

where *E*_*k*_ is the DFT of the even-indexed part of *f*_*m*_, and *O*_*k*_ is the DFT of the odd-indexed part of *f*_*m*_. Based on the periodicity of the DFT:
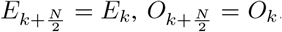. We have the following:

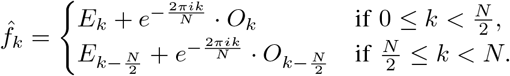

The harmonic factor *e*^−2*πik/N*^ have the property that:

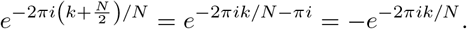

Thus, we can cut the number of harmonic factor calculations in half. For 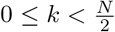:

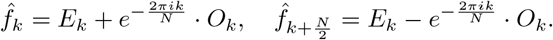

The pseudocode for Cooley-Tukey FFT algorithm is detailed in Algorithm 1. The time complexity of FFT is *O*(*N* log *N*).

##### Fast Fourier Transform 2D

is an extension of FFT 1D. First, we have the DFT 2D as:

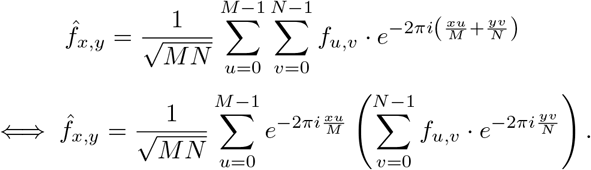

We can compute 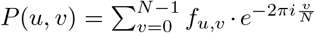 by FFT 1D. Again, we use FFT 1D to compute

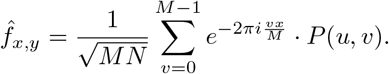

###### Algorithm 1

Cooley–Tukey FFT algorithm

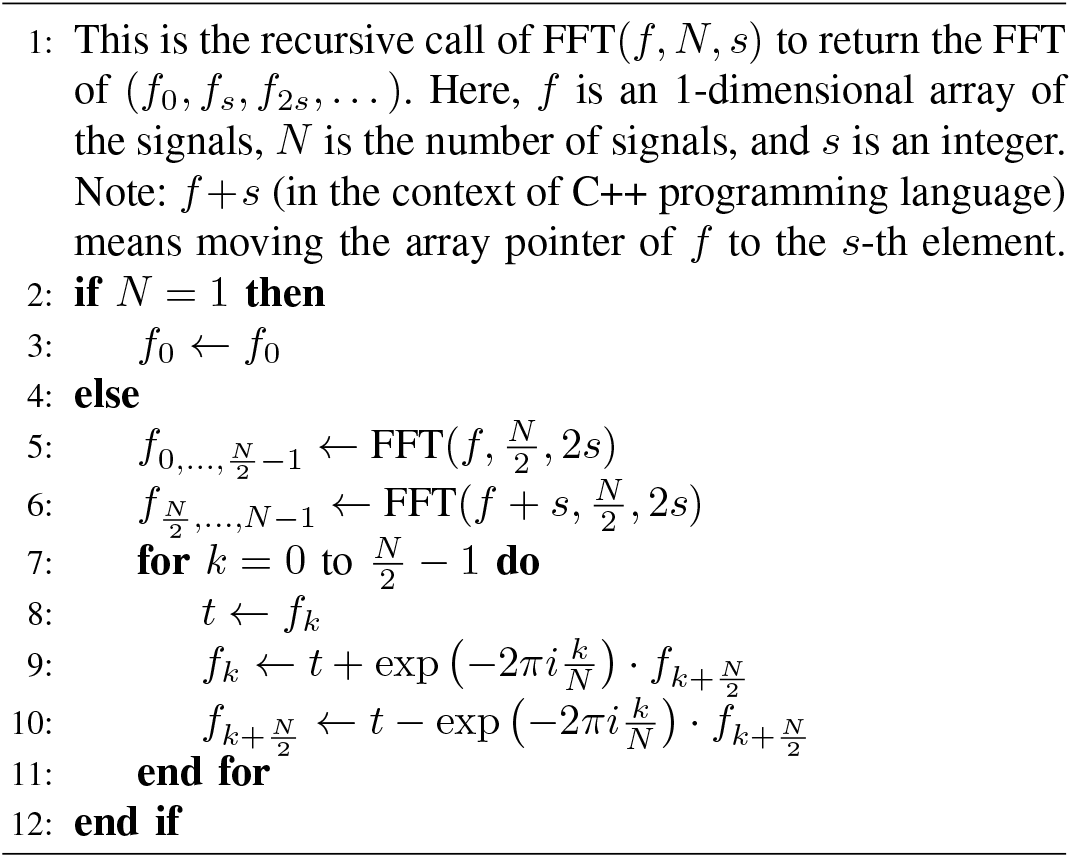

On the another hand, the inverse pass:

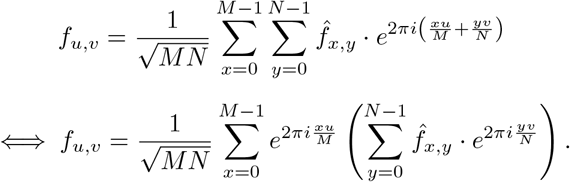

We compute 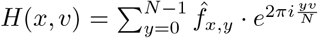 and then 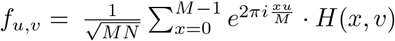 by FFT 1D algorithm.

#### B. Architecture

##### a) Complex ReLU

The network leverages the ReLU activation function to introduce nonlinearity and promote sparsity. The ReLU function transforms all negative entries in the matrix to zero while preserving the positive values. The complex ReLU function adheres to the Cauchy-Riemann condition when the real and imaginary parts are both strictly positive or negative. Complex ReLU is defined as the sum of the ReLU-applied real and imaginary components. Mathematically, it is expressed as follows:

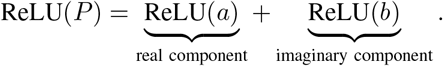

##### b) Complex BatchNorm2d

Batch Normalization [69] (BN) is used to adjust the input value distribution of each layer in a neural network to match a standard normal distribution, characterized by a mean of 0 and a standard deviation of 1. For complex numbers, it is not straightforward to apply BN because translating and scaling them to have a mean of 0 and variance of 1 leads to complications. Specifically, the variances of the real and imaginary parts are not normalized equally, leading to a possibly highly eccentric elliptical distribution. Thus, the data distribution changes when viewed as a two-dimensional vector. Given a batch input *x*, the normalized 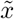 is calculated using the mean and variance as follows:

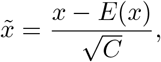

where *C* represents the covariance matrix and *E*(*x*) is the mean of the data. The matrix *C* is a 2 *×* 2 matrix defined as follows: where *R*(*x*) and *I*(*x*) are the real and imaginary parts of *I*(*x*), respectively. To normalize the distribution of features to be learned by the network, learnable reconstruction parameters are incorporated, analogous to normalization in the real number domain. The shift parameter, however, is more complex, having two learnable components (both real and imaginary). The scaling parameter is a 2 *×* 2 positive semi-definite matrix with three degrees of freedom. The complex BN process includes three learnable components. The complex BN can be mathematically described as:

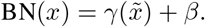

